# Predictive masking is associated with a system-wide reconfiguration of neural populations in the human visual cortex

**DOI:** 10.1101/758094

**Authors:** Joana Carvalho, Remco J. Renken, Frans W. Cornelissen

**Affiliations:** Laboratory of Experimental Ophthalmology, University Medical Center Groningen, University of Groningen, Groningen, the Netherlands; Cognitive Neuroscience Center, University Medical Center Groningen, University of Groningen, Groningen, the Netherlands

**Keywords:** prediction, masking, population receptive field, connective field, reorganization, artificial scotoma, fMRI

## Abstract

The human visual system masks the perceptual consequences of retinal or cortical lesion-induced scotomas by predicting what is missing from nearby regions of the visual field. To reveal the neural mechanisms underlying this remarkable capacity, known as predictive masking, we used fMRI and neural modeling to track changes in cortical population receptive fields (pRFs) and connectivity in response to the introduction of an artificial scotoma (AS). Consistent with predictive masking, we found that extrastriate areas increased their sampling of the V1 region outside the AS projection zone. Moreover, throughout the visual field and hierarchy, pRFs shifted their preferred position towards the AS border. A gain field model, centered at this border, accounted for these shifts, especially for extrastriate areas. This suggests that a system-wide reconfiguration of neural populations in response to a change in visual input is guided by extrastriate signals and underlies the predictive masking of scotomas.

## Introduction

When the information extracted from a visual scene is incomplete, the visual system attempts to predict what is missing based on information from nearby regions of the visual field. A remarkable perceptual consequence is the masking of retinal lesions, which makes patients remain unaware of their partial loss of vision. Consequently, such masking often results in delayed diagnosis and treatment (1, 2) of such lesions. The underlying process to which we will refer to as predictive masking (PM), also plays a prominent role in healthy perception, e.g evident from the masking of the blind spot, the receptorless area of the retina where the optic nerve leaves the eye, and from many visual illusions in which color, brightness, or textures spread into and mask neighbouring regions of the visual field (3, 4). Consequently, the process is sometimes also popularly referred to by this behavioral manifestation as “filling-in”.

Despite the scientific and clinical relevance of PM, its underlying neuronal mechanisms are still poorly understood. Human and animal physiology studies into PM and studies of the neural consequences of retinal lesions have shown receptive field (RF) expansion and shifts in RF preferred position towards spared portions of the visual field (5–9). However, such RF changes also occur following simulated scotomas, thus suggesting that these changes may not result from structural plasticity (10–12). Indeed, the observed RF changes may be an indirect consequence of a modulation in the responses of neurons in the scotoma projection zone (SPZ), possibly caused by gain adjustments that reduce the feedforward information (13–16), a downregulation of inhibition (17), or a change in feedback from higher order areas with large RFs (18–21).

Such observations have led to the controversial hypothesis that predictive masking is explained by neurons modifying their receptive field properties, (22) while the precise neural basis of PM remains unknown. In addition, previous studies assumed that PM is a local process restricted to the SPZ, so they focused on the SPZ and the early visual cortex. However, if PM is a consequence of functional changes (changes in gain), we would expect neurophysiological modifications to occur both inside and outside the SPZ and throughout the visual hierarchy. In the present study, we therefore tested the hypothesis that PM involves a global reconfiguration of RFs and their connectivity. Specifically, in analogy to the behavioral phenomenon, we expect that in the cortical region responsible for PM, the neural mechanisms within the SPZ should show a decreased reliance on information from within the SPZ and an increased reliance on the information from outside of it. If this hypothesis is confirmed, we could create more accurate models of visual perception and improve diagnostic methods for patients with visual field defects.

To test our hypothesis, we used functional MRI in combination with biologically-inspired neural population modeling to track changes in RF properties and cortical connectivity following the introduction of an artificial scotoma (AS) into the visual field of human participants (thus mimicking a lesion to their visual system). We modeled the observed changes in pRF preferred position using a gain field model and we examined how cortical connections between recording sites (connective field size) changed in response to the AS.

## Results

Retinotopic mapping was performed under three different stimulus conditions: a conventional retinotopy stimulus based on luminance contrast (LCR) used for delineating visual areas, an artificial scotoma stimulus (AS^+^) and a control stimulus identical to AS^+^ but without the artificial scotoma (AS^-^). The stimuli used in the two AS conditions were designed to stimulate the Low Spatial Frequency (LSF) selective neurons predominantly. The LSF carries coarse information about the visual scene and it is presumably encoded mainly by neurons with large RFs (23, 24). This is expected to facilitate PM. The AS^-^ and AS^+^ conditions were used to define the pRFs size and preferred position (PP) for each voxel (see materials and methods section for additional details).

### The scotoma border attracts pRFs

To examine the presence of changes in pRF properties between the AS^-^ and AS^+^ conditions, the data for the four different quadrants (each containing one AS) was collapsed onto a single quadrant. Next, the pRF properties of the voxels were spatially binned based on their preferred position (PP) as estimated in the AS^-^ condition. In visual area V1, following the presentation of an AS, pRFs with a PP originally inside the AS shifted radially outwards and towards the border of the AS (Figure 1A). However, an analysis of the entire V1 representation showed that pRFs outside of the AS also appear to be attracted towards the AS (Figure 1B). These shifts were observed across the visual hierarchy (Figure S1 and S2). We compared the PP in both conditions across the visual hierarchy using a two-way repeated measures ANOVA, which revealed main effects of condition (AS^-^ versus AS^+^, F(1,35)=8.4, p=0.004) and ROI (F(5,35) = 4.09, p= 0.003). Furthermore the PP shifts were more pronounced for extrastriate areas (the interaction between ROI and condition was significant (F(5,35)=7.87, p=0.0034), see Figure S1). *Post hoc* tests (FDR corrected) showed significant differences in position between conditions for all the visual areas tested (p<0.001). These observations suggest that pRFs throughout the visual field shifted their PP towards the AS border. When analyzed in more detail, Figure 1C shows how the PP shifted as a function of the pRFs’ distance to the center of the AS. Note that the shift is minimal at the border (at 2.5 deg.). Figure D plots the radial component of PP shift, again as a function of the pRFs distance to the AS center. This shows a nearly perfect linear relationship between the radial shift and the pRFs’ initial PP (r^2^< −0.99 and p<1×10^-8^ for all the visual areas, Figure S2). Note that pRFs situated at the AS border hardly shift radially. Additional analyses excluded that these patterns are simply the result of statistical or modeling biases (Figures S3 and S4).

**Figure 1.**
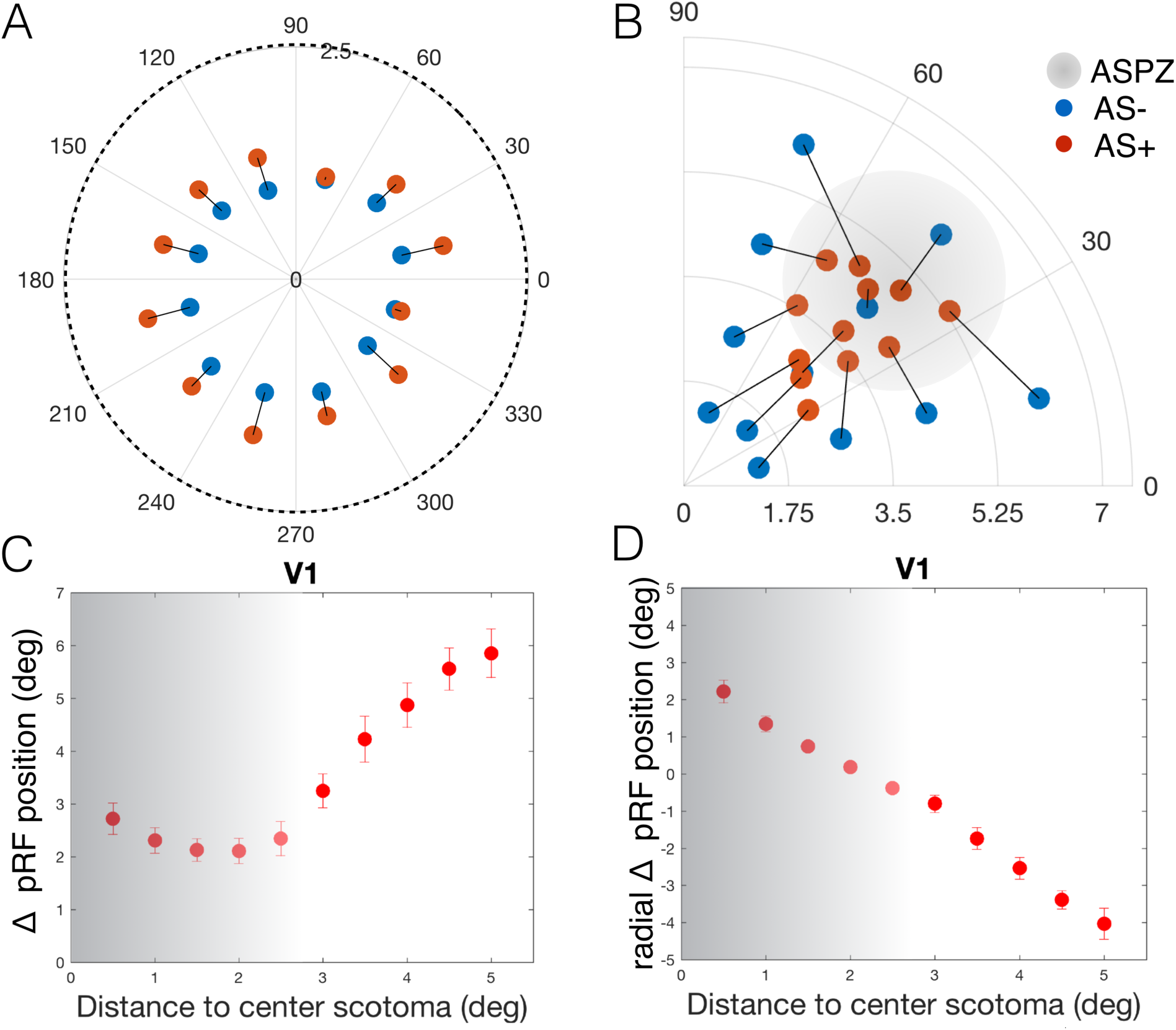
V1 pRF position change in response to AS. A: Shift between the two conditions AS^-^ (blue) and AS^+^ (red) of the pRFs with initial PPs located inside the ASPZ, averaged across participants. B: Position change between conditions in various sectors of the visual field, averaged across participants. C: pRF position change (AS^+^ vs AS^-^) as a function of distance between pRF position (based on AS^-^) and the center of the scotoma (bins of 0.5 deg., Euclidean space). Error bars show the standard error of the mean over hemispheres. D: pRF position change projected onto the radius as a function of the radial distance between pRF position measured in the AS^-^ and the center of the scotoma. The gray transparent region refers to the AS, the darker region corresponds to the center if the AS. Figure S2 shows the results for the other visual areas, V2-LO2. Figure S3 shows that these results are not simply due to random position noise. The AS^+^ results were obtained using the Scotoma Field (SF) model (which minimizes model biases). The pRF position shifts between AS^-^ and AS^+^ were present using either model (SF and FF, Figure S4).

### A gain field model explained the artificial scotoma induced pRF position shifts

The systematic changes in pRF PP suggest that these shifts may depend on their position relative to the AS border. Such shifts can be modeled using a gain field (GF) (25). To determine whether the border plays a critical role in the pRF reconfiguration, we first plotted the radial component of the shifts (Figure 2A). This indicates that the shifts are of similar magnitude all around the perimeter of the AS (although different for pRFs initially inside or outside the AS). Next, we determined if we could predict the radial component in the AS^+^ condition based on the PPs in the AS^-^ condition by modulating the AS effect using a GF that is centered on the AS border (Figure 5B). Figure 2 shows the predicted and measured pRF positions shifts (Panel 2B) and size ratios (Panel 2C). The GF model performed well and explained 50% and 92% of the variance in the radial position shifts and size changes, respectively (Figures 2B and C). Figure 2D shows that the position predictions of the GF model are most accurate for the higher order areas (V1, VE =39%; LO1, VE=66%). The PP shifts tend to increase along the visual hierarchy (Figure S1). Although the pRF sizes increased with eccentricity and visual hierarchy (Pearson’s correlation coefficient: r^2^>0.8 and p<0.05 for all the visual areas tested), the pRF PP change does not strongly correlate with the pRF size within every visual field map (V1 r^2^= 0.06; V2 r^2^=-0.06; V3 r^2^= 0.13; V4 r^2^= −0.06; LO1 r^2^= 0.1; LO2 r^2^=-0.2; all p<0.0005). Regarding changes in the pRF size, a comparison across condition and visual areas revealed that the pRF size does not change significantly between conditions (F(1,35)=0.007, p=0.93) but it does change with visual area (F(5,35)=6.5, p<0.0001), and the interaction between condition and visual area is not significant (F(5,35)=0.63, p=0.67). *Post hoc* tests (FDR corrected) did not show any significant differences in pRF size between all the conditions tested p>0.09).

**Figure 2.**
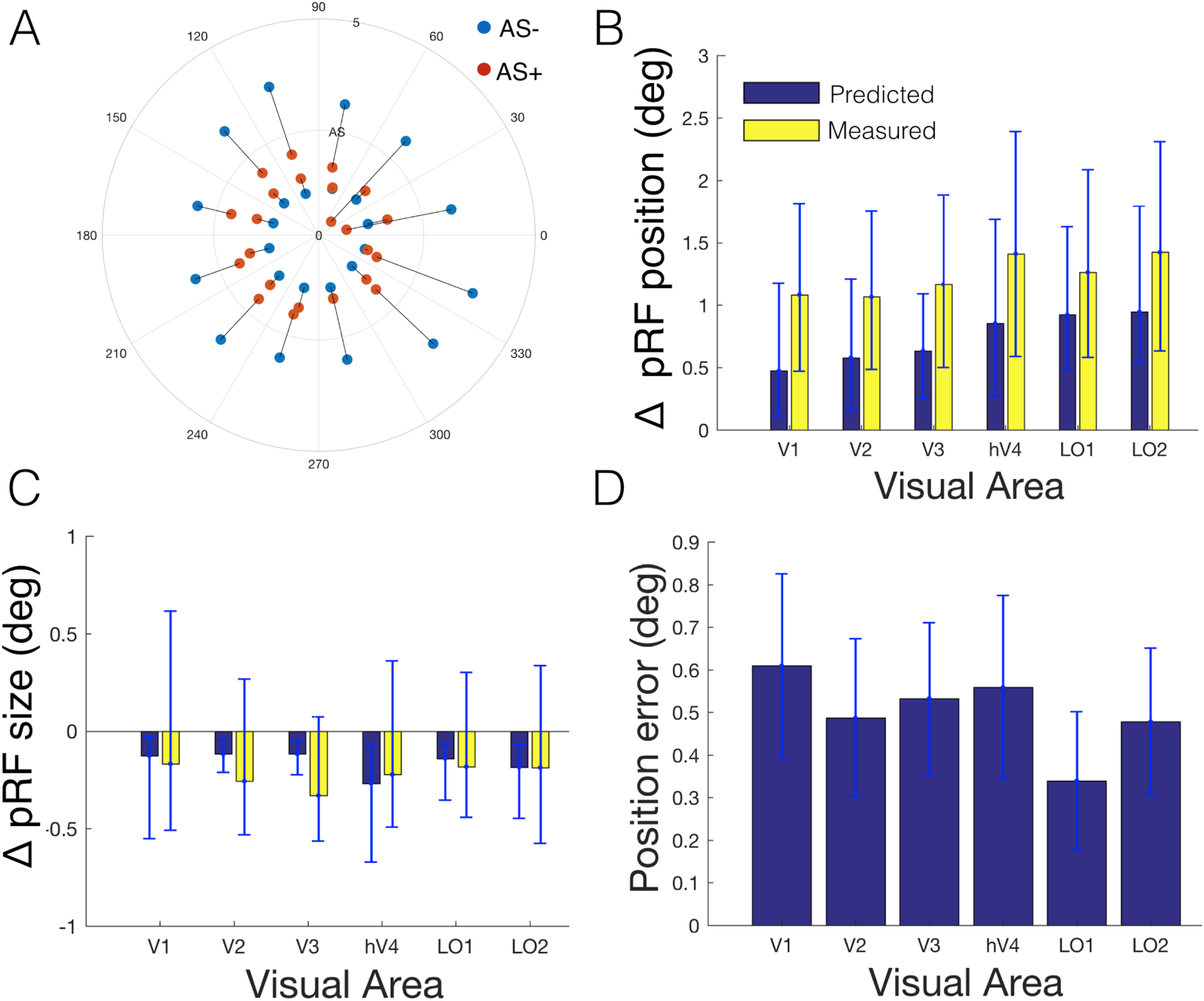
A gain field model centered at the AS border explains changes in preferred population RF position. A: Radial position change between the two conditions AS^-^ (blue) and AS^+^ (red) in various sectors of the visual field inside and outside the AS, averaged across participants. The region inside the AS corresponds to the ASPZ. B/C: Measured (yellow) and predicted pRF position shifts (B) and size changes (C) in response to an AS. D: Mean average error between the predicted and measured pRF shifts. The error bars represent the interquartile ranges across the voxels in the test set. The estimated GF size did not vary significantly between visual areas (F=0.16; p=0.97)).

### Neural populations in extrastriate cortex increase their V1 sampling region

Visual areas beyond V1 may also respond to the AS by changing their V1 sampling. Changes in sampling of a source area such as V1 can be quantified by modeling the connective field (CF) of the recording site. The CF enables the prediction of the neuronal activity of a recording site (voxel) in a target region (e.g. V2) given the activity in another part of the brain (e.g. V1). CFs are estimated without modeling the stimulus, so they are not subject to modeling bias and may reflect other components of brain function, such as feedback signals. Changes in the CFs may thus arise independently from the V1 pRF changes reported above. Figure 3A shows the difference in CF size between the two AS conditions (AS^+^ - AS^-^) for the voxels whose PP was initially located either inside or outside the ASPZ. For some visual areas (voxels initially inside the ASPZ) the CF became larger following the introduction of the AS. In particular, LO1 recording sites inside the ASPZ sampled from a larger region of V1, which is evident from the increased CF size. This effect was not clearly present for recording sites outside the ASPZ (LO1: inside ASPZ, p=0.002; outside ASPZ p=0.14). To show how the accumulation of these changes influences the sampling of V1, we projected the CFs back into visual field space by convolving them with the V1 pRFs from which they sample. To isolate AS-induced changes in the CFs from those in the pRFs of V1, the CFs of both the AS^-^ and AS^+^ conditions were back projected using the same set of pRFs (those from the AS^-^ condition). For areas V2 and LO1, Figure 3B shows the CF sampling density in the conditions AS^-^ and AS^+^ and their difference (AS^+^ - AS^-^). Overall, V2 sampling density is reduced in the AS^+^ compared to the AS^-^ condition. This effect is most pronounced for recording sites within the ASPZ. The ΔCF image shows that the introduction of the AS generally resulted in a denser sampling of V1 regions outside the ASPZ. This effect seems particularly pronounced in LO1.

**Figure 3.**
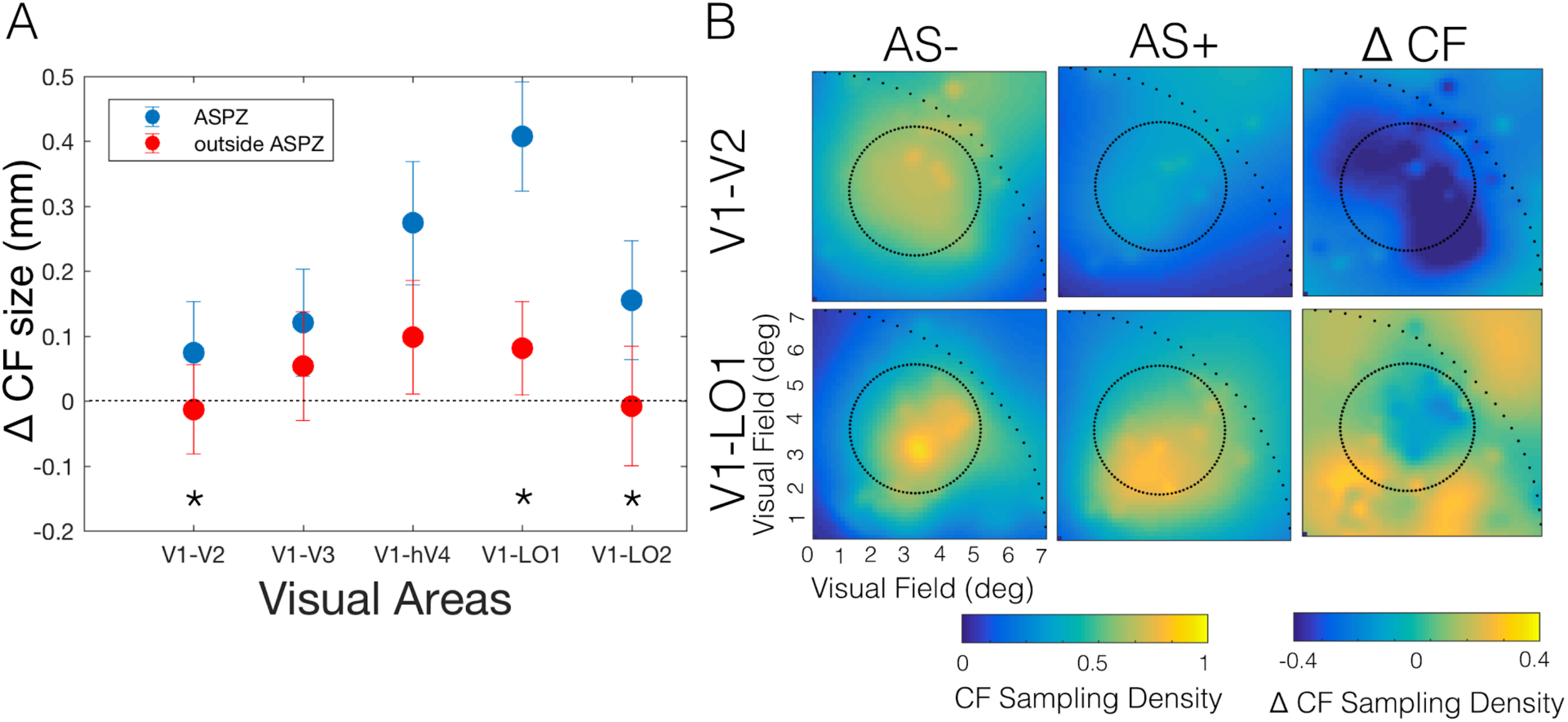
Changes in cortico-cortical connections in response to AS. A: Difference in the cumulative distribution at the 90% point (dashed line Figure S5A and B) of the CF size between AS^+^ and AS^-^ in (blue) and outside (red) the ASPZ. The error bars represent the 5% and 95% CI. The CF sizes between the two conditions (AS^-^ and AS^+^) differed significantly inside and outside scotoma for V2, LO1 and LO2 (p<0.001), represented in the graph by *. Figures S5 A and B show the cumulative histogram of the CF sampling extent for ASPZ of the visual areas tested. Note that the V1 sampling extent increases (shift to the right) with visual hierarchy. This trend is not present for the voxels located outside the ASPZ (Figure S5B). The significance level between the two conditions (AS^+^and AS^-^) per ROI is shown on the bottom right of the cumulative graphs. B: Coverage map of CFs obtained for AS^-^, AS^+^ and the difference between the two conditions (AS^+^ - AS^-^). The back projection onto the visual field was performed based on the pRF estimates obtained with AS^-^. The sparse dotted line depicts the visual stimulation area and the dotted line the AS location in the visual field. Each map represents the combined data from 7 subjects.

## Discussion

Our main finding is that in extrastriate cortical regions, in particular LO1, we observed increased sampling of V1 regions outside the ASPZ, which would be required for the predictive masking of the scotoma. Moreover, we find that inside and outside the ASPZ and throughout the visual hierarchy, pRFs reconfigured their preferred spatial position and shifted it towards the AS border. This behavior is inconsistent with what would be expected based on PM. However, a gain field model, centered at the AS border, could effectively explain these changes. This suggests that the pRF changes primarily serve to focus neural resources on regions of potential interest and constitute a component of normal visual perception. The model explained the shifts most effectively for extrastriate areas, in particular for area LO1. We therefore postulate that the population modifications originate in extrastriate areas and, through feedback, also modulate the V1 pRFs. Therefore, changes in intra-area connectivity (connective fields), rather than those of the pRFs, may be the neural underpinning of PM. In summary, our results reveal an extended, system-wide reconfiguration of neural population properties in response to the change in visual input evoked by an AS. Below, we discuss our findings and interpretation in detail.

### Extrastriate cortex increased its sampling of V1 outside of the ASPZ

To understand how the cortico-cortical connections between visual areas change in response to an AS, we quantified their CFs, which describe how extrastriate target areas (V2 to LO2) sample from source area V1. Dissociating the changes in the extrastriate CFs from their pRF shifts revealed an increased sampling density of the V1 region outside of the ASPZ in response to the AS. This effect was particularly evident for LO1, where the sampling from V1 increased especially for voxels inside the ASPZ. This indicates that cortico-cortical connections change following the presentation of the AS, resulting in increased capturing of information from outside the scotomatic region. This is consistent with PM.

The capacity to dissociate connectivity from the visual input via the back projection of the CFs is less susceptible to stimulus-related model-fitting biases (due to its independence from the stimulus) and informs how the visual information is integrated across different cortical areas. It also has the potential to capture the neural circuits underlying pRF dynamics (Carvalho et al., in press).

### Feedback from extrastriate regions drives system-wide reconfiguration

Previous studies have reported dissociation in the representation of superficial, middle and deep layers of V1. In these studies, the superficial and deep layers represented the feedback mechanisms that modulate perception, and the middle layers represented the visual input. Evidence of predictive feedback in the superficial layers of V1 was found when neurons were deprived of information in a partial occlusion paradigm (26, 27). Selective feedback-associated activation of the deep layers of V1 was also found in a study on the Kanizsa illusion (28). Therefore, the pRF changes measured in the early visual cortex could plausibly be driven by feedback connections from extrastriate cortex. Moreover, based on our results and those of others, extrastriate area LO1 is a potential candidate for the origin of these feedback signals. It plays a major role in the processing of oriented boundaries or borders (29, 30) and its role can be dissociated from that of LO2, which preferably processes shape (30). In our analysis, the gain field model best explained the observed pRF modulations in this area, which would be expected for signals originating in this area. Moreover, the increased sampling of V1 was most prominent for LO1 voxels. We therefore propose that the reconfiguration of neural populations in response to an AS is modulated by extrastriate signals and may underlie predictive masking.

Although PM is linked to perceptual filling-in (FI), we opted to not quantify perceptual FI during our experiments. This is because such a perceptual task could interfere with the attention task and increase the chance of unintentional small eye-movements in the direction the AS, thereby actually decreasing FI. Therefore, we performed psychophysical tests outside the scanner and prior to the present study, which indicated that participants reach stable FI after about 30 sec (Figure S6). Since our actual mapping experiment started after 60 sec and the design of the retinotopic stimulus was optimized to yield FI, we assumed that observers were filling in the AS at the time we performed the pRF and CF mapping.

### An artificial scotoma induces a system-wide reconfiguration of neural population receptive fields

In response to the AS, pRFs shift their preferred spatial position towards the AS border. While such shifts have been reported previously for pRFs initially located inside the natural SPZ (31) and ASPZ (10–12, 32), our study is the first to show that this reconfiguration is not restricted to the ASPZ, but is a system-wide phenomenon. Within the ASPZ, the pRFs shifted their preferred position towards the AS border, which could be consistent with an extrapolation process. Following the shift, pRFs are more likely to be activated by spared portions of the visual field, and can thus contribute to the spatial masking of the scotoma. However, the pRFs initially located outside the ASPZ shifted their preferred position towards the AS border as well. These pRFs are more likely activated by non-stimulated portions of the visual field. Therefore, this behavior cannot easily be reconciled with PM.

Previous studies have suggested that changes in the pRF properties in response to an AS can result from a model bias driven by partial stimulation of the neuronal populations (11, 12, 32). This effect can be controlled by taking into account the presence of the AS during the pRF modeling (12, 32). Accordingly, we used two pRF modeling approaches: one that assumed the presence of the AS – the Scotoma Field (SF) model, and one that did not – the Full Field (FF) model. We found similar positional shifts with both models, thus indicating that our findings are unbiased (Figure S4). Importantly, CFs are not affected by such model biases.

### A gain field at the scotoma border explains the shifts in pRF preferred position

The factor common to all shifts is that these were predominantly directed towards the AS border. Indeed, the PP changes could be explained by a biologically motivated GF that accounts for the presence of the AS. This suggests that the presence of an AS results in a reweighting of the spatial response selectivity towards the scotomatic border. Similar results were found using a model of attention (25, 33). Therefore, the presence of the AS could result in a deployment of attention towards the AS border. Although the AS was designed to induce PM (filling-in), a reduced visual stimulation may actually be salient to the early visual system (34). In this case, the PP shifts indicate that the border was a salient feature. This interpretation is supported by the fact that GF model described the PP shifts accurately, especially for the extrastriate areas. This interpretation is also in line with previous studies, which showed that high-level mechanisms (attention) modulate perception via feedback projections (20). The reconfiguration of neural population properties may therefore have the more general role of allocating neural resources to salient features in the visual field. This may help to scrutinize these in more detail, or alternatively, to resolve prediction errors (35).

This interpretation links to previous hypotheses about the underlying mechanisms of PM, in particular the suggestion that the masking of an AS results from (slow - tenths of seconds) adaptation to salient features (such as a border) in combination with a fast extrapolation process (36). Although, the design of the present experiment did not allow us to separate these two components, the GF model can shed some light on these issues. We suggest that during the border adaptation, neural resources are allocated to the borders of the scotoma in response to its saliency, resulting in a reconfiguration of the RFs and consequently in the predictive spatial masking of the scotoma. These findings indicate that the modulation of the pRF structure by cognitive factors contributes to the adaptation to the scotoma borders and consequently to the predictive masking.

In contrast to previous studies using retinal and cortical scotomas (5, 8), our observed PP shifts were not accompanied by increases in pRF size (if something they tended to shrink). The absence of size changes in early visual cortex may be related to our use of a low spatial frequency stimulus. Therefore, the most responsive neurons defining the pRF already had large receptive fields, leaving little room for further expansion. Importantly, the presence of the AS did not alter fundamental structural characteristics of the visual cortex, such as the increase of the pRF size over eccentricity and visual hierarchy. However this last aspect does not explain the increase of the position shifts over the visual hierarchy.

### Limitations and future studies

Eye movements may bias pRF estimates and commonly result in increased pRF sizes (25, 38, 39). Eye movements were not recorded during scanning but were minimized by having observers perform an attention task that demanded central fixation. Moreover, eye movement artifacts should have resulted in increased pRF sizes, which we did not find.

For five of the seven observers the AS^-^ and AS^+^ conditions were performed in two different scan sessions raising the possibility that pRF shifts were due to misalignment between the functional and anatomical scans. However, such shifts should all have been in the same direction, e.g. fovea to periphery. Moreover, we find similar shifts in the two observers who performed the two conditions within the same scan session. Therefore, we conclude that the observed pRF shifts are genuine.

We defined the pRFs contained by the ASPZ based on the pRF estimates obtained with the AS^-^ condition. As an alternative method, we also defined the ASPZ based on a scotoma localizer in which the AS and its background were stimulated separately. The results obtained using either definition of the ASPZ resulted in highly analogous findings, reason why we choose to present the results based on only one method.

Future studies measuring the neuronal mechanisms associated with PM at finer scale (e.g. at higher fMRI resolution) could reveal changes that are masked at a coarser scale. This is not only because one can identify more pRFs in the ASPZ, but also because it enables determining laminar profiles across cortical depth, which could help to determine at which level of cortical processing the feedback and feedforward signals modulate perception.

In conclusion, in the present study we have shown that partial occlusion of local visual input results in a system-wide reconfiguration of the RF properties of neural populations and their connectivity. Furthermore, we suggest that this reconfiguration is guided by extrastriate signals, that the reconfiguation is an integral component of normal perception and that it forms the basis of predictive masking in health and disease.

## Materials and Methods

### Participants and Ethics statement

Seven participants (3 females; average age: 28; age-range: 26–32) with normal or corrected-to-normal vision were included in the study. The participants indicated that they understood the instructions. Prior to participation, participants signed an informed consent form. Our study was approved by the Medical Ethical Review Board of the University Medical Center of Groningen, and conducted in accordance with the Declaration of Helsinki.

### Data acquisition

Stimuli were presented on an MR compatible display screen (BOLDscreen 24 LCD; Cambridge Research Systems, Cambridge, UK). The screen was located at the head-end of the MRI scanner. Participants viewed the screen through a tilted mirror attached to the head coil. Distance from the participant’s eyes to the display (measured through the mirror) was 120 cm. Screen size was 22×14 deg. The maximum stimulus radius was 7 deg of visual angle. Visual stimuli were created using MATLAB (Mathworks, Natick, MA, USA) and the Psychtoolbox (40, 41).

### Stimuli

#### Luminance-contrast defined retinotopy (LCR)

LCR consists of a drifting bar aperture defined by high-contrast flickering texture (42). The bar aperture, i.e. alternating rows of high-contrast luminance checks drifting in opposite directions, moved in 8 different directions (four bar orientations: horizontal, vertical and the two diagonal orientations), with two opposite drift directions for each orientation (Figure 4A). The bar moved across the screen in 16 equally spaced steps each lasting 1 TR. The bar contrast, width and spatial frequency were 100%, 1.75 and 0.5 cycles per degree, respectively. After 24 steps (one pass and a half), 12 s of a blank full screen stimulus at mean luminance was presented.

**Figure 4.**
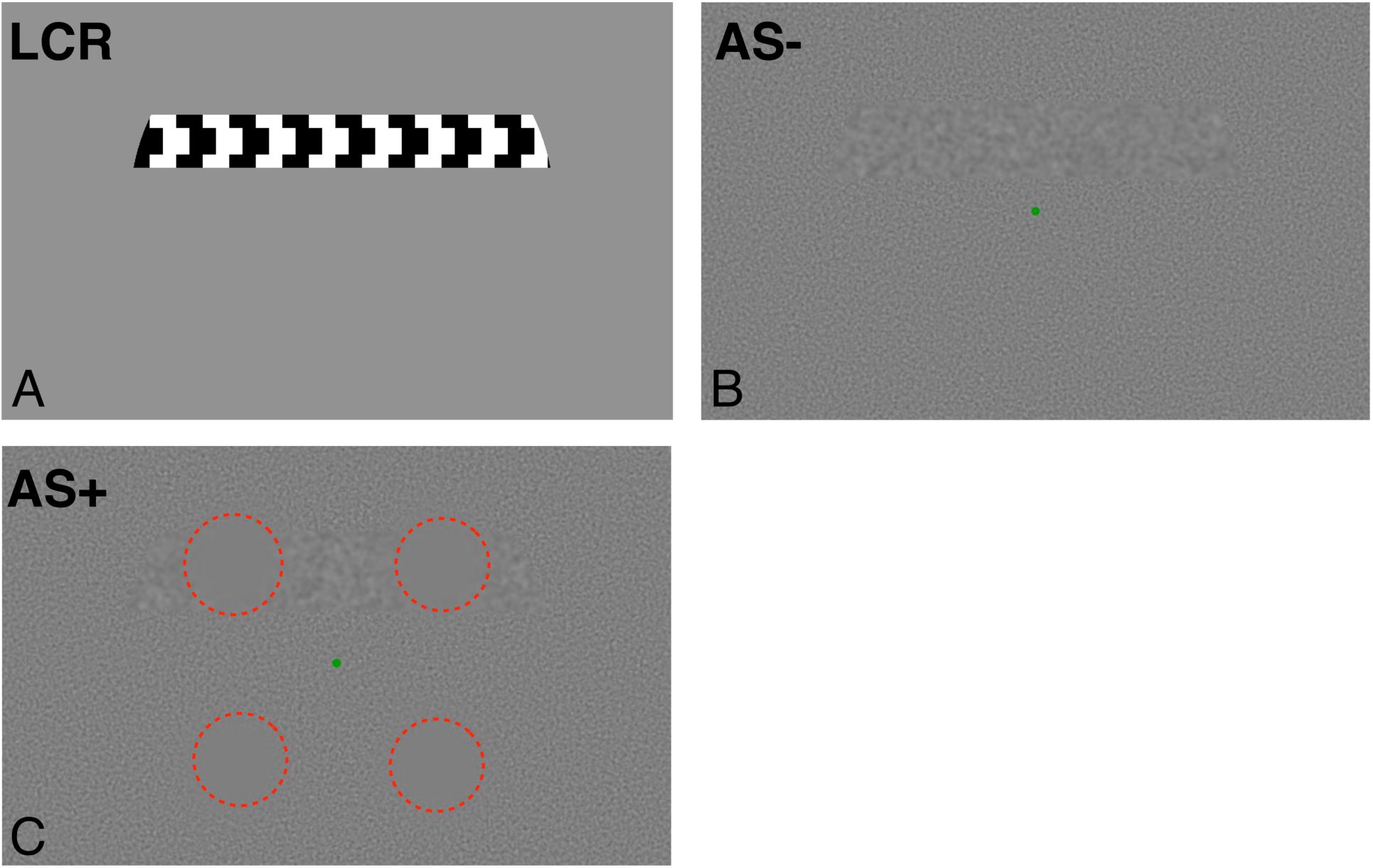
Example of the stimuli used to obtain pRF parameter estimates. A: LCR; B: AS^-^, C: AS^+^, for visualization purposes the AS are outlined with a dashed red line. NB: this red dashed line was not presented to the participants

#### Artificial Scotoma (AS) conditions

The stimuli used in the two AS conditions were adapted from the LCR stimulus. More specifically, the bar and background could be distinguished from each other only on the basis of their spatial frequency (Figure 4B). The AS^-^ condition served as the control condition for the AS^+^ condition that contained the actual scotoma. The bar’s movement directions and orientations matched those of the LCR condition. The width of the bar aperture was 3 degrees. The bar content was dynamic white-noise band passed filtered at frequencies from 0 to 2 cycles per degree (cpd). The background consisted of dynamic white SF band passed from 2 to 4 cpd. The long edges of the bar were smoothed using an exponential mask. The formula for this mask was: 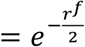, where r is the distance to the center-line of the bar, and *f* the mask factor. The value of *f* was fixed at 4. The bar moved at a speed of 0.46 deg/sec. The AS-condition was used to define a baseline PP and size of the pRF for each voxel. The AS^+^ condition was similar to AS^-^ (with equal bar aperture size, movement and SF). Four ASs were superimposed on the dynamic noise background (see Figure 4C). The scotomas were centered at each quarter field at 4.5 deg of eccentricity. Each AS consisted of 2.5 deg radius disc tapered by an exponential mask at the edges, similar to the masking of the bar: 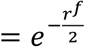, where, r is the distance from the center of the scotoma and f is fixed at a value of four, as before. Preceding each run was a one-minute adaptation period during which the participants viewed only the background with the AS superimposed while performing the fixation attentional task. In psychophysical experiments, performed prior to the fMRI scans, we determined that this period was sufficient to induce filling-in (see Figure S6).

### Attentional task

During scanning, participants were required to perform a fixation task in which they had to press a button each time the fixation point turned from green to red. The average performance on this task was above 86% for all the conditions. The task performance per condition is shown in Table S1.

### MRI scanning and preprocessing

Scanning was carried out on a 3 Tesla Siemens Prisma MR-scanner using a 64-channel receiving head coil. A T1-weighted scan (voxel size, 1mm^3^; matrix size, 256 x 256 x 256) covering the whole brain was recorded to chart each participant’s cortical anatomy. The functional scans were collected using standard EPI sequence (TR, 1500 ms; TE, 30 ms; voxel size, 3mm^3^, flip angle 80; matrix size, 84 x 84 x 24). Slices were oriented to be approximately parallel to the calcarine sulcus. For the retinotopic scans LCR and AS^-^ a single run consisted of 136 functional images (duration of 204 s) and for AS^+^ a single run consisted on 168 functional images (252 s).

The T1-weighted whole-brain anatomical images were re-sampled to a 1 mm^3^ resolution. The resulting anatomical image was automatically segmented using Freesurfer (43) and subsequently edited manually. The cortical surface was reconstructed at the gray/white matter boundary and rendered as a smoothed 3D mesh (44).

The functional scans were analyzed in the mrVista software package for MATLAB (available at http://white.stanford.edu/software). Head movements between and within functional scans were corrected (45). The functional scans were averaged and co-registered to the anatomical scan (45), and interpolated to a 1mm isotropic resolution. Drift correction was performed by detrending the BOLD time series with a discrete cosine transform filter with a cutoff frequency of 0.001Hz. To avoid possible saturation effects, initial images were discarded for the LCR and AS^-^ (8 TRs), as well as for the AS^+^ (40 TRs). Note that the full 60 seconds adaptation period was removed for the AS^+^.

### Experimental procedure

Each participant completed two fMRI sessions of approximately 1.5 h. In the first fMRI session, 5 participants were subjected to the anatomical scan and LCR, and they performed the AS^-^ experiment (6 runs, 3.4 min each). In the second fMRI session, the AS^+^ experiment (6 runs, 4.2 min each) were performed. To eliminate the possibility that differences between conditions (AS + and AS-) would result from the acquisition in different sessions, these were performed for 2 participants (S06 and S07) in the same session.

### Visual field mapping: pRF modeling

The pRF analysis was performed using both conventional pRF mapping (42) and a custom implementation of the Monte Carlo Markov Chain (MCMC) Bayesian pRF approach (46, 47). In the conventional method, a 2D-gaussian model was fitted with parameters: center (x0, y0) and size (σ - width of the Gaussian) for each voxel. All the parameter units are in degrees of visual angle and are stimulus-referred. We used SPM’s canonical Haemodynamic Response Function (HRF) model. The conventional pRF estimation was performed using the mrVista (VISTASOFT) Matlab toolbox. The Bayesian pRF approach enables the estimation of the uncertainty associated with each pRF parameter. The uncertainty was defined by the 25% and 75% quantiles of the estimated distribution.

In both approaches, the data was thresholded by retaining the pRF models that explained at least 15% of the variance. Furthermore, the functional responses to LCR, AS^-^ and AS^+^ were analyzed using the FF model. The AS^+^ condition was also analyzed using the SF model (Figure 5A).

**Figure 2.**
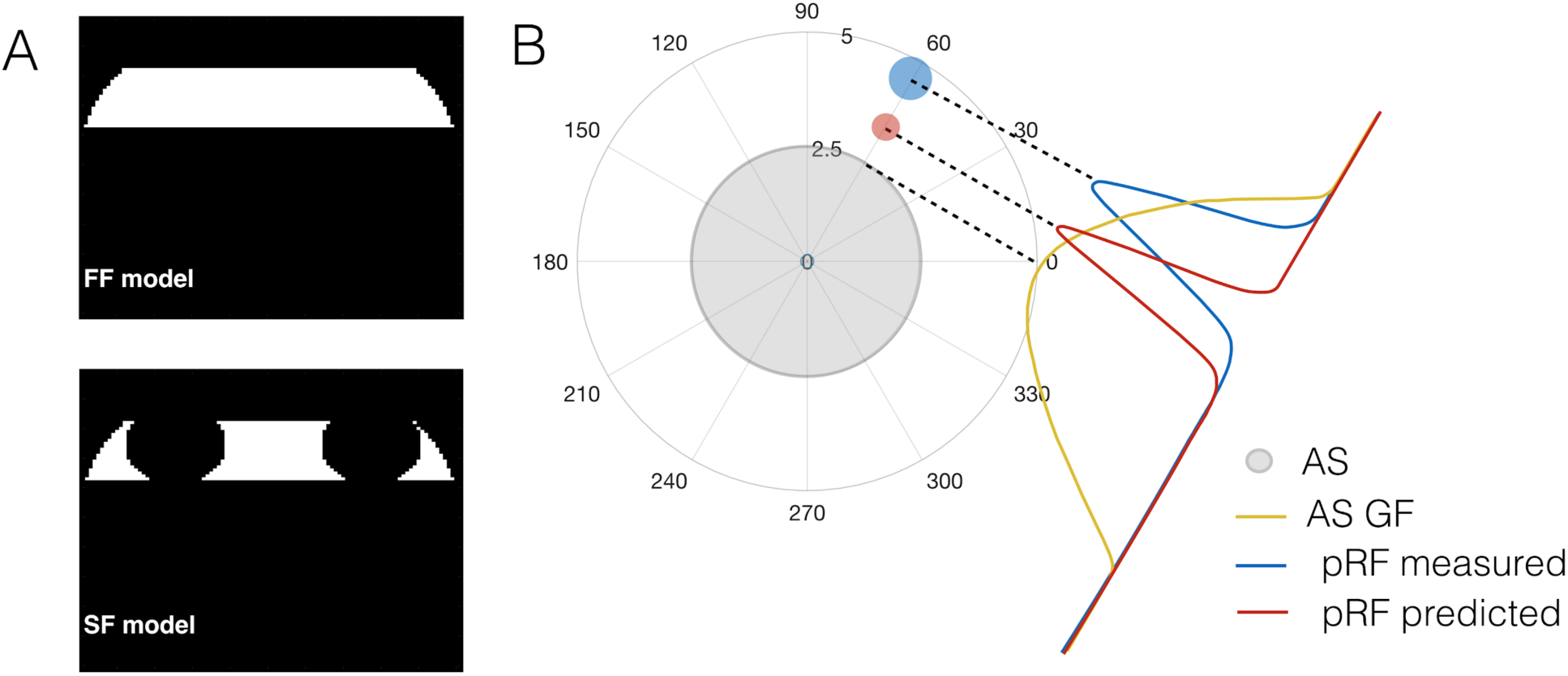
Models of neural responses used in the analysis, FF, SF and AS Gain Field model. A: The full field (FF) and scotoma field (SF) models used in the pRF analysis. B: AS GF model: the AS (shaded grey region) effect was modeled as the AS GF (yellow), centered at the edge of the scotoma closest to the pRF (blue). This results in a predicted pRF (red), shifted towards the scotoma.

### ROI and Artificial Scotoma Projection Zones definition

The cortical borders of visual areas were derived based on phase reversal, obtained with the conventional pRF model using the classical the LCR stimulus. Per observer, six visual areas (V1, V2, V3, V4, LO1 and LO2) were manually delineated on the inflated cortical surface.

Based on the pRF estimates obtained with the AS^-^ condition, the ASPZ was defined as the voxels for which the pRF was completely contained within the AS regions of the visual field.

### Gain Field model

The influence of the AS on the pRF’s preferred position and size was modeled as a gain field (GF), i.e., the multiplication of two Gaussian components (25, 33, 37, 48). In our study, the first Gaussian component corresponded to the pRF estimated in the AS^-^ condition (*u_AS_*_−_,*σ_AS_*_−_). The second Gaussian component corresponded to the GF (*u_GF_*,*σ_GF_*) elicited by the AS: it represented the influence of the AS on the pRF’s preferred position. The GF was centered on the border of the AS at the point nearest to the original pRF location (Figure 5). The product of these two components resulted in a third Gaussian (*u_pAS+_*,*σ_pAS+_*), that represented the predicted pRF in the AS^+^ condition. Equations 1 and 2 show how the properties of the third Gaussian were derived.

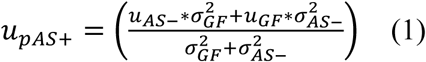

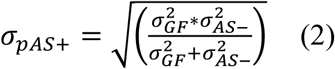

The GF size was estimated by minimizing the error between the predicted and the measured position shifts, which is the radial distance between the AS^+^ and AS^-^. For verification of the model’s accuracy, the data was split into a training set (50% of the data) and a test set (the remaining 50% of the data).

### Connective field (CF) modeling

The CF model predicts the neuronal activity of a recording site (voxel) in a target region (e.g. in V2) given the aggregate activity in a source region (in V1) (49). The fMRI response of each voxel is predicted using a 2D circular Gaussian CF model, folded to follow the cortical surface of the source region. The CF output parameters are the position and spread (size) across the source surface. Given a CF position and a size, a time-series prediction is then calculated by weighting the CF with the BOLD time series. The optimal CF parameters are found by minimizing the residual sum of squares between the predicted and the measured time-series. In this study, only CFs with a VE> 0.6 were retained.

### Statistical analysis

Data was thresholded by retaining the pRF models that explained at least 15% of the variance in the BOLD response in the three conditions (LCR, AS^+^, AS^-^). For the analysis of changes in pRF properties in response to the AS, the pRF estimates of the four quadrants were collapsed onto a single quadrant. Subsequently, voxels were binned into 12 bins, each covering an eccentricity range of 1.75 deg and a polar angle range of 30° (Figure 3B). Additionally within the ASPZ, voxels were binned into 12 bins of 30 deg of polar angle each after shifting the origin to the center of the ASPZ (Figure 3A).

The PP change corresponds to the Euclidean or radial distance between the AS^+^ and AS^-^ conditions. The size ratio, σ*_r_*, was calculated based on the following equation:

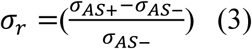

The CF coverage maps were obtained by back projecting each CF into the visual space using the pRFs for V1 obtained with AS^-^. First, per voxel in the target region, a CF was calculated, i.e. the target voxel is expressed as the weighted (CF factor) average of the signals measured in V1 (the source region). As the pRF was known for each voxel in V1, we calculated the spatial sampling by summing all pRFs of V1 weighted by the CF factor. The total CF coverage map was calculated by summing these maps across all voxels in the target region. Finally, a group average (n=7) was calculated across subjects.

Repeated measures ANOVA, with ROI, condition (AS^-^, AS^+^SF), hemisphere and position bin as within-subject parameters, was used to compare the difference of the pRF preferred position and size between conditions. Subjects were treated as random variables. For the AS^+^ condition, the pRF properties were estimated using two different models (FF, SF Figure 2A). Separate statistical analyses were performed for each of the resulting parameter sets. Permutation tests (1000 replications) were used to determine significance level of the differences in CF size between conditions inside and outside the ASPZ. For this, data was aggregated over participants and condition labels were permuted.

All analyses were performed using MATLAB (version 2016b; Mathworks, Natick, MA, USA) and R (version 2.11.1; R Foundation for Statistical Computing, Vienna, Austria). A p-value of 0.05 or less was considered significant.

## Acknowledgments

Author JC was supported by the European Union’s Horizon 2020 research and innovation programme under the Marie Sklodowska-Curie grant agreement No. 641805 (NextGenVis) awarded to FWC. The funding organization had no role in the design, conduct, analysis, or publication of this research.

## Supplementary material

### 1. Shifts in PP of pRFs occur throughout the visual hierarchy

**Figure S1.**
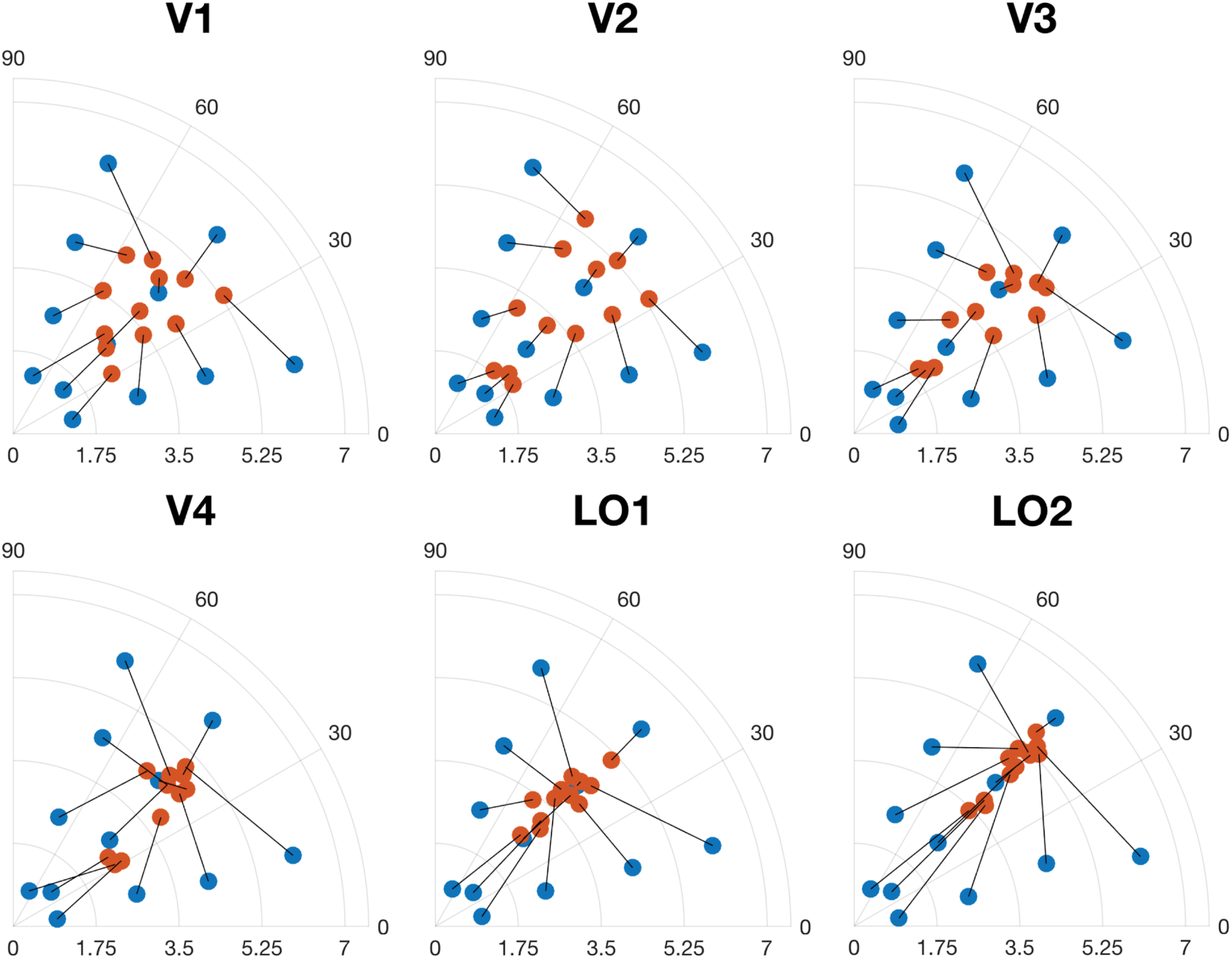
Changes in pRF position in response to the presentation of an AS. The figures show position changes between the AS^-^ and AS^+^ conditions in different sectors of the visual field, averaged across participants. The V1 data is the same as shown in figure 1.

**Figure S2.**
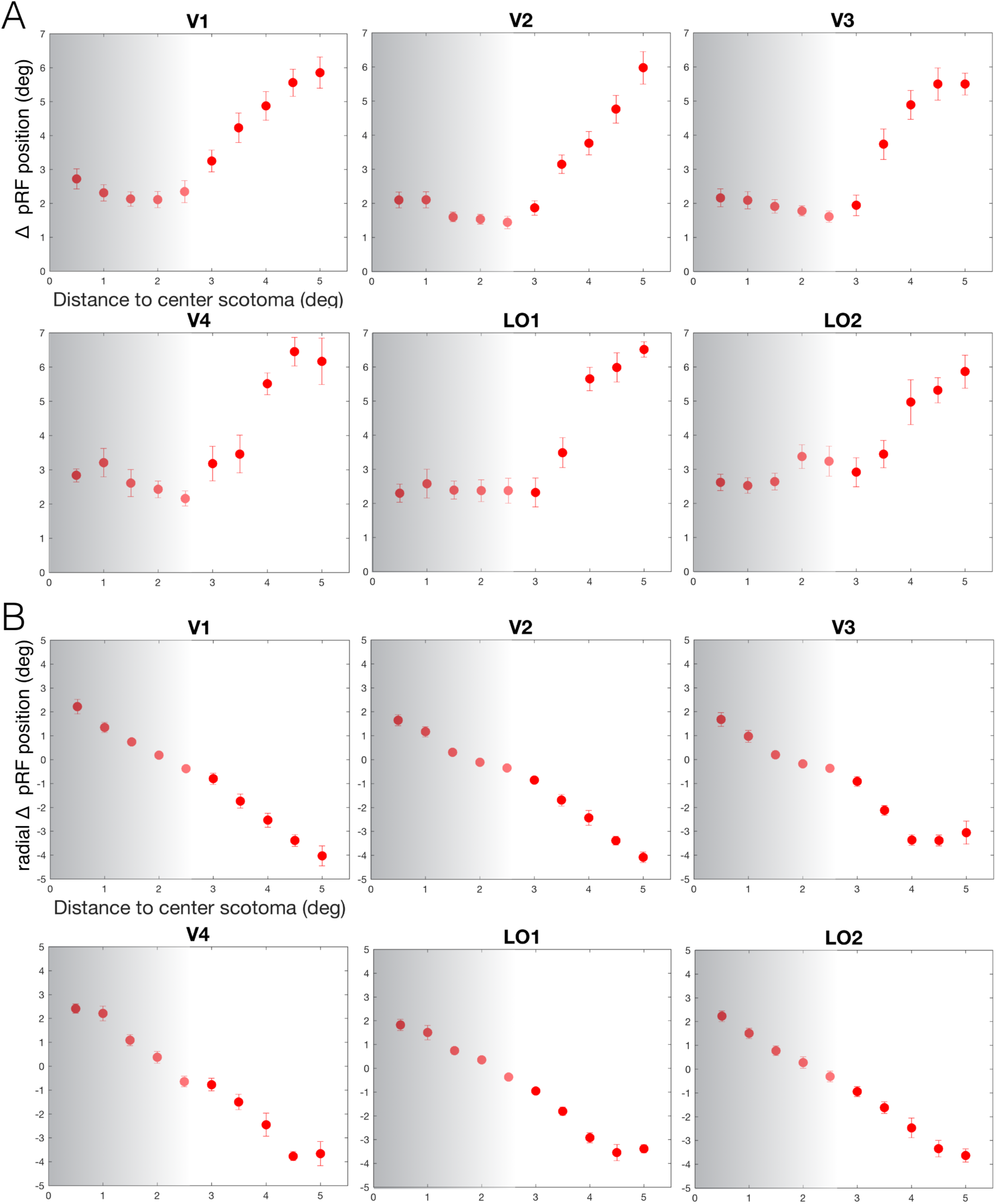
Change in preferred position of pRFs as a function of their distance to the AS. A: pRF position change as a function of the Euclidean distance between the pRF position and the center of the scotoma (bins of 0.5 deg) for all the visual areas analysed. The error bars represent the standard error. B: Change in radial pRF position as a function of the radial distance between the pRF position and the center of the scotoma.

### 2. Simulations of pRF shiftss

To verify that pRF shifts did not result from a statistical bias (regression to the mean), we simulated the Euclidean and radial pRF position change resulting from arbitrary shifts in position. We simulated 10000 pRF positions uniformly distributed across the stimulated visual field for both conditions (AS^+^ and AS^-^). PRF’s PP were collapsed onto a single quadrant and the Euclidean and radial PP shifts were binned in 0.5 degree bins as a function of the distance to the center of the AS. Figure S3 shows a comparison between the simulated and measured pRF PP shifts. For both types of shift (radial and Euclidean) the observed shifts cannot be explained as a result of a statistical bias. Note that in panel B at the edge of the scotoma (2.5 deg) the measured position shift is ∼ 0 deg whereas the simulated shift is ∼1.5 deg. Moreover the voxels located near the center of the scotoma are displaced of 2.2 deg (corresponding to distance between the center of the AS to its edge) while the simulated displacement is the double.

**Figure S3.**
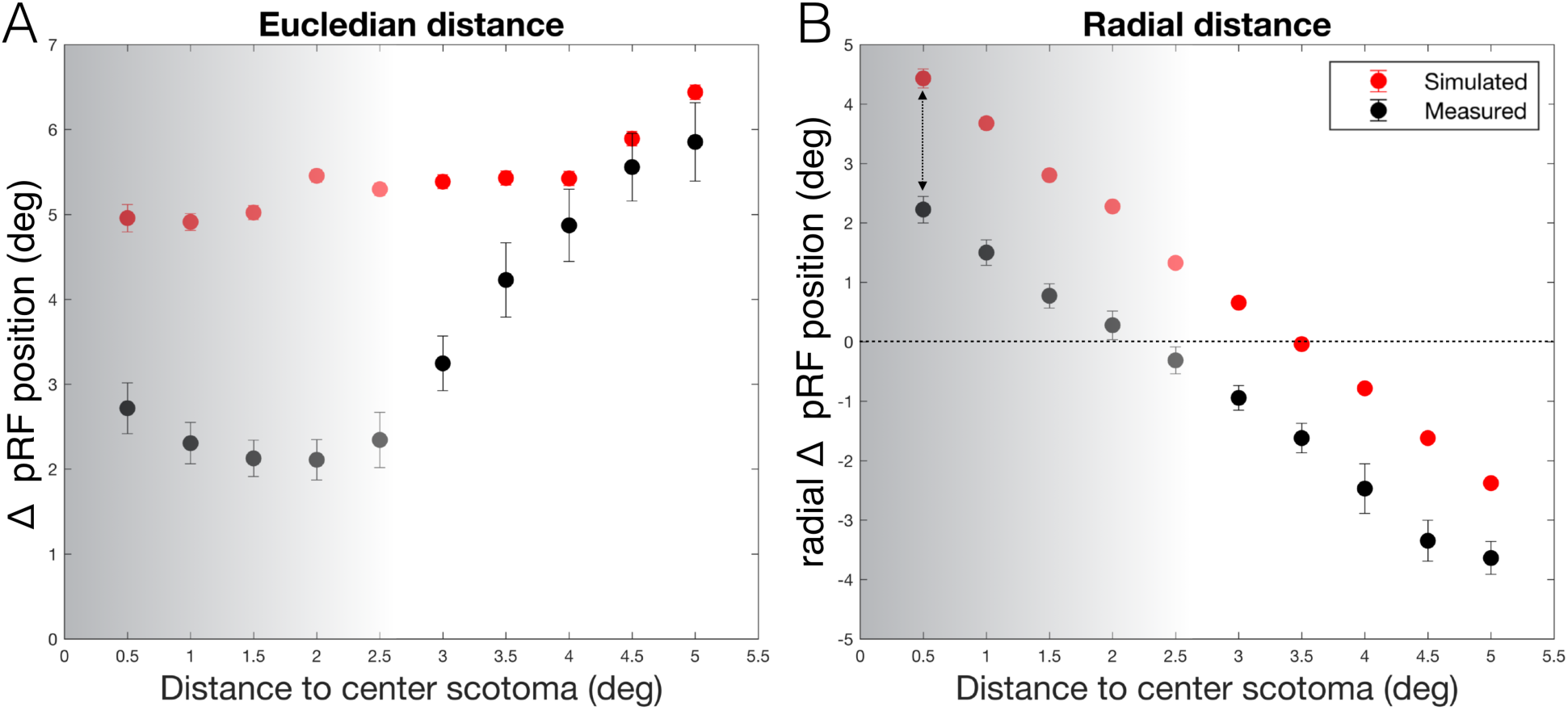
Simulated position change as a function of the distance to the AS. A: Simulated pRF position change as a function of the Euclidean distance between pRF position measured with AS^-^ and the center of the scotoma, in bins of 0.5 deg. Error bars show the standard error of the mean over hemispheres. B: pRF position change as a function of the radial distance between pRF position measured with AS^-^ and the center of the scotoma.

### 3. Comparison between SF and FF model analyses

Previous work has suggested that pRF shifts may result from disregarding the AS when creating a model of the stimulus input that drives the pRF. In the main body of our paper, we described a model that took the AS into account (scotoma field (SF)). Here, we show the effect of using a full field (FF) model. The pRF position shifts between AS^-^ and AS^+^ conditions were present when applying either of the both models. Furthermore, the presence of the artificial scotomas neither reduced the BOLD amplitude nor affected the explained variance of the models.

**Figure S4.**
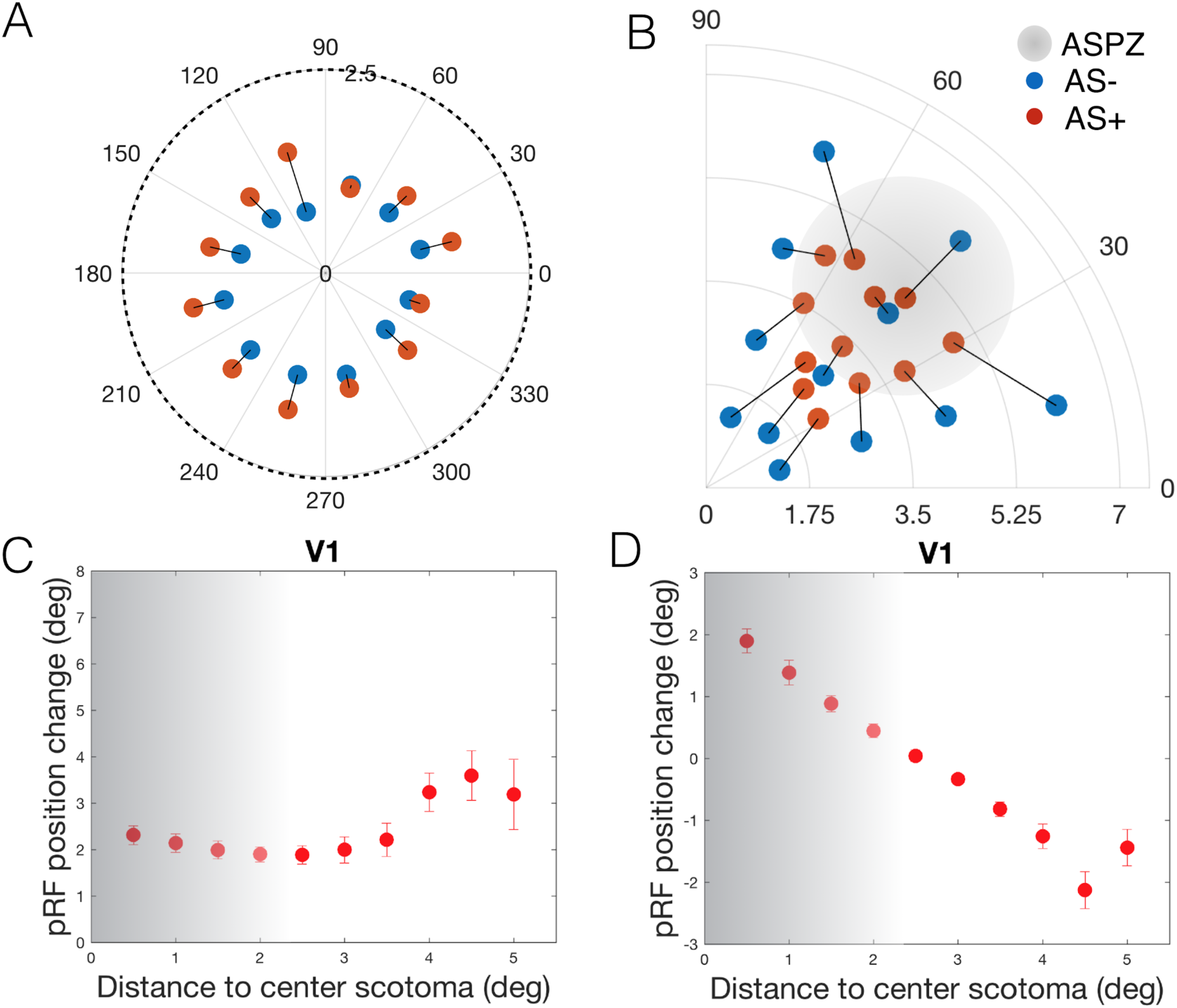
Changes in V1 pRF position change in response to the presentation of the AS as calculated when using a full field (FF) model. A: Shift between the two conditions AS^-^ (blue) and AS^+^ (red) of the pRFs with initial PPs located inside the ASPZ. B: Position change between conditions in different sectors of the visual field, averaged across participants. C: pRF position change (AS^+^ vs AS^-^) as a function of distance between pRF position (based on AS^-^) and the center of the scotoma (bins of 0.5 deg, Euclidean space). Error bars show the standard error of the mean across hemispheres. D: The change in radially projected pRF position change as a function of the radial distance between pRF position measured in the AS^-^ and the center of the scotoma. The gray transparent region refers to the AS, the darker region corresponds to the center of the AS.

### 4. Connective fields in extrastriate cortex increase their sampling extent

**Figure S5.**
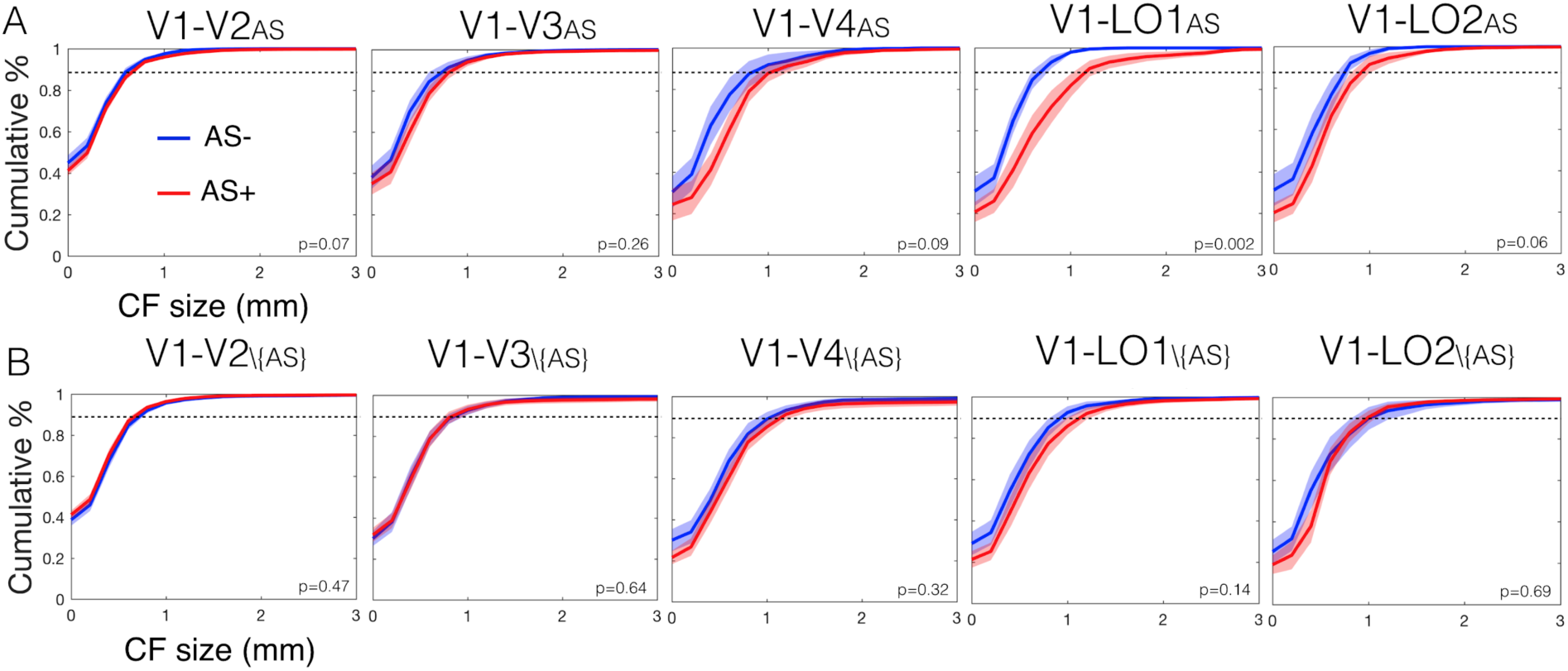
CF changes in response to AS. A: Cumulative percentage of the CF size for the conditions AS^-^ (blue) and AS^+^ (red) calculated for the voxels within the ASPZ in the target visual areas V2 a LO2. B: Analogous analysis to panel A, but for those voxels outside the ASPZ. The shaded area represents the 5% and 95% confidence intervals. The p-value on the bottom right of each graph shows the significance of the difference between the two conditions.

### 5. Filling-in time

Six of the seven participants included in the MRI study, participated in a psychophysical experiment to establish the time required for filling-in to occur. The stimulus consisted of dynamic white noise band pass filtered at frequencies of 2 to 4 cpd. Four AS with a radius of 2.5 deg were superimposed. The participant’s task was to fixate in the center of the screen (represented by a white dot – 0.15 deg radius) and press a button when the background was perceived as uniform (the AS had been filled in). Filling-in time corresponded to the time interval since the presentations of the scotomas until the button press was recorded. The scotomas were centered at 4.5 deg eccentricity, at each quarter field. Per participant four repetitions (trials) were performed. Between two consecutive trials there was a gap of 15s during which a uniform grey background was shown in order to prevent carryover. The filling-in time was always less than one minute (figure S6). Therefore one minute of filling-in time was allowed in the fMRI experiment for all participants.

**Figure S6.**
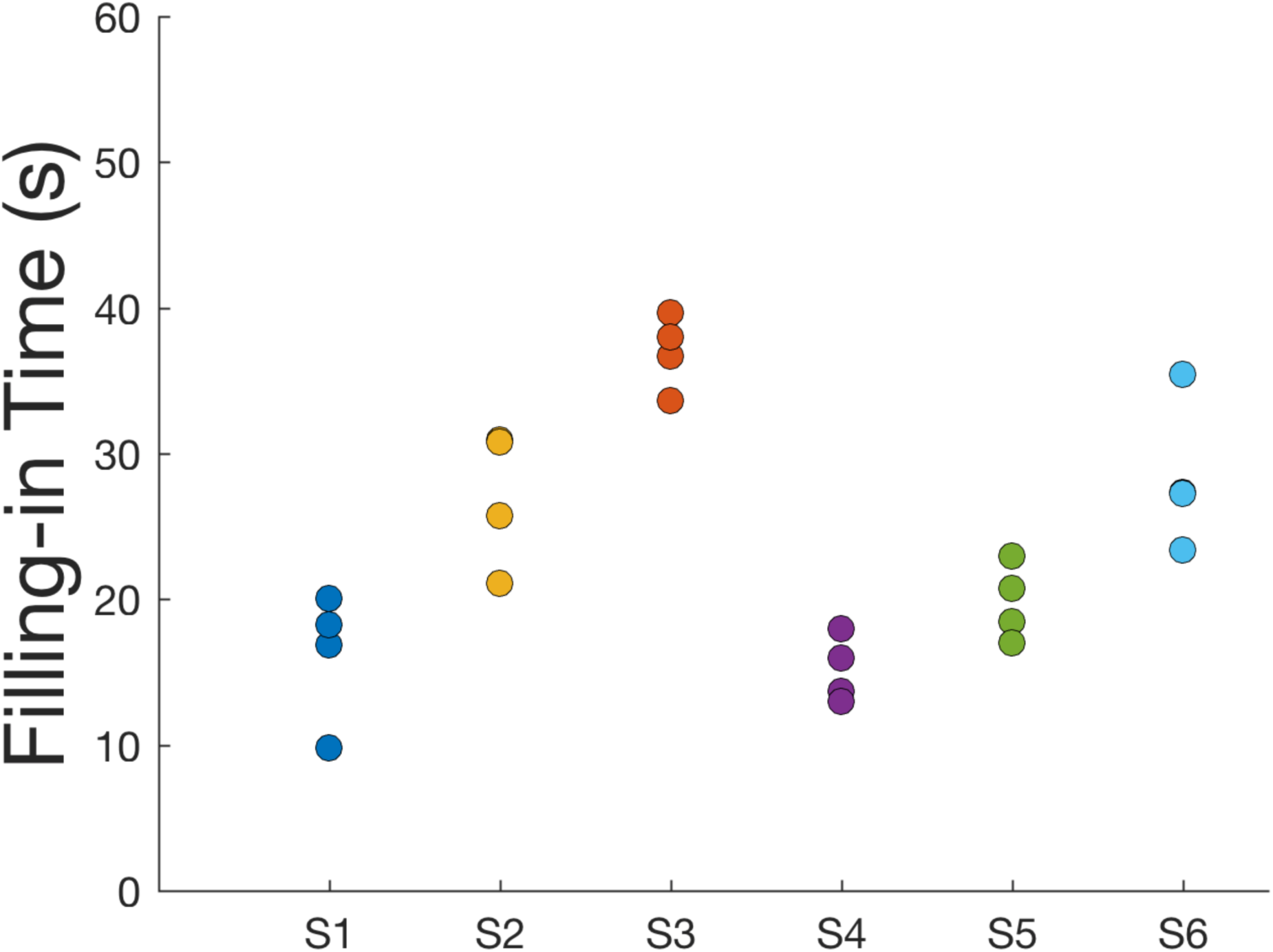
Results of the psychophysical tests used to define the optimal stimulus parameters (adaptation time). Filling-in time measured per trial and per participant.

### 6. Attention task performance

**Table S1.** Performance (average and standard error) of the attention task per condition. One-way repeated measures ANOVA showed no significant difference between the attention task performance between the conditions AS^+^ and AS^-^ (p=0.6341).

| Task | Mean (%) | Standard error (%) |
| --- | --- | --- |
| LCR | 90.9 | 6.8 |
| AS- | 86.0 | 8.7 |
| AS+ | 87.7 | 3.4 |

## Notes

Conflict of Interest: None of the authors has potential conflicts of interest to be disclosed.

